# Live imaging-assisted domain-specific CRISPR genome editing at single cell resolution in plants

**DOI:** 10.1101/793240

**Authors:** Ting Li, An Yan, Elliot M. Meyerowitz

## Abstract

CRISPR (Clustered Regularly Interspaced Short Palindromic Repeats) technology has been widely used for genome engineering in a wide range of organisms^1^, but much of the development of CRISPR-based genome editing has been aimed toward improving its efficiency and accuracy, so as to obtain genetic materials carrying known and stably heritable genome modifications. Precise spatiotemporal control over genome editing technology at cell type resolution is a key challenge for gene function studies. Some tissue-specific CRISPR genome editing methods relying on phenotypic characterization and fluorescent immune-staining techniques have been developed for biomedical research and gene therapy, they function by spatially controlling expression of Cas9 ^2^. Recent work establishes the presence and location of mutational events at a single cell level in *Arabidopsis* roots and stomata^3,4^. Here we present an efficient domain-specific CRISPR-Cas9 system combined with a high resolution live-imaging based screening strategy, applied in the shoot apical meristem of *Arabidopsis thaliana*. Using the system we investigate PIN-FORMED1 (PIN1) protein functions in tissue morphogenesis and PIN1 mechanical stress response in a cell layer-specific fashion. We find that reported failure to generate new primordia in epidermal PIN1 knockout SAMs is due to a reduction in mechanical stress differences in the sub-epidermal layer. The methods described are applicable to spatial-temporal gene manipulation in plants.

## Main

The initial test of the live imaging-assisted CRISPR system was to knock out a GFP reporter sequence in a promoter expression domain-specific manner, allowing genome editing efficiency to be examined by live imaging of GFP fluorescence *in vivo*. To achieve high potential efficiency, we developed a CRISPR system to express Cas9 protein and sgRNA at high levels ^5^. This was done by driving the expression of CAS9 protein^6^ mainly in the epidermis of *Arabidopsis* plants using the promoter of the *MERISTEM LAYER 1 (ATML1)* gene^7^ and driving expression of guide RNA (sgRNA) targeting a sequence in the *GFP* gene with four different *Arabidopsis* RNA Polymerase III promoters (*AtU6-26, AtU6-1, AtU3b* and *AtU3d*), all driving the same sgRNA (Fig. S1a, b)^5,8,9^. Although we utilized multiple RNA Polymerase III expression cassettes to overexpress gRNA in an attempt to increase the gene modification efficiency, side by side comparison is necessary to confirm that this was effective. When estimating the gRNA binding specificity (http://crispr.mit.edu), none of 14 listed potential off-target sequences (score ranges from 0.0% to 0.8%) is within an exon.

As an initial proof of principle, we examined the action of the expression domain-specific CRISPR in an *Arabidopsis* nuclear *pATML1:H2B-mGFP* reporter line^10^, which shows strong epidermal expression, and also weaker subepidermal expression of a histone protein tagged with GFP in the shoot apical meristem (Fig. 1a, d, Fig. S4a-c). Screening with live imaging, we found that 19 out of 90 independent T_1_ transgenic GFP CRISPR founder plants showing diminished epidermal marker expression in the SAM (Table S1). No visible developmental defect was observed in mutated lines. In most cases, the mutation pattern was homogeneous, in that GFP signal is not detected in any cells in the epidermis of the SAM (Fig. 1b, e). In one mutation line, a few cells in the epidermis of a flower primordium of the SAM showed GFP signal, suggesting a mosaic mutational pattern within the targeted gene expression domain (Fig. 1c, f). The GFP signal is specifically suppressed only in the first layer (L1) and GFP signal in cells of the second layer (L2) of the mutated SAM remained intact, indicating specificity of mutagenesis to cells where Cas9 is expressed at high levels (Fig. 1m-o). Since AtML1 gene expression is not detected in plant sperm cells and egg cells^11,12^, and the epidermal layer is not thought to be the source of germ cells of plants^13,14^, finding mixed siblings of T_2_ plants with wild type-like (9/15, 60%, Fig. 1g-i) and knockout signal patterns (6/15, 40%, Fig. 1j-l, Table S1) indicates that inheritance of the CRISPR constructs renews the domain-specific mutagenesis effect in each generation, making the original lines a source of continued mosaicism for analysis. PCR-based single clone Sanger sequencing from dissected shoot apex tissue detected a certain percentage (4-12%) of CRISPR-induced somatic mutations in the genomic DNA from 5 out of 6 T_2_ founder lines that showed L1-specific GFP signal suppression, confirming that the mutations happened within the gRNA binding site, while no mutation was detected in wild type genomic DNA samples (Table S2). Moreover, distinct mutational genotypes were detected in two axillary meristems from the same mutation line (line #4-1, #4-2, Table S2) or in the same SAM (line #2, #4-1, #6, Table S2), indicating mosaicism in this CRISPR system.

**Figure 1.**
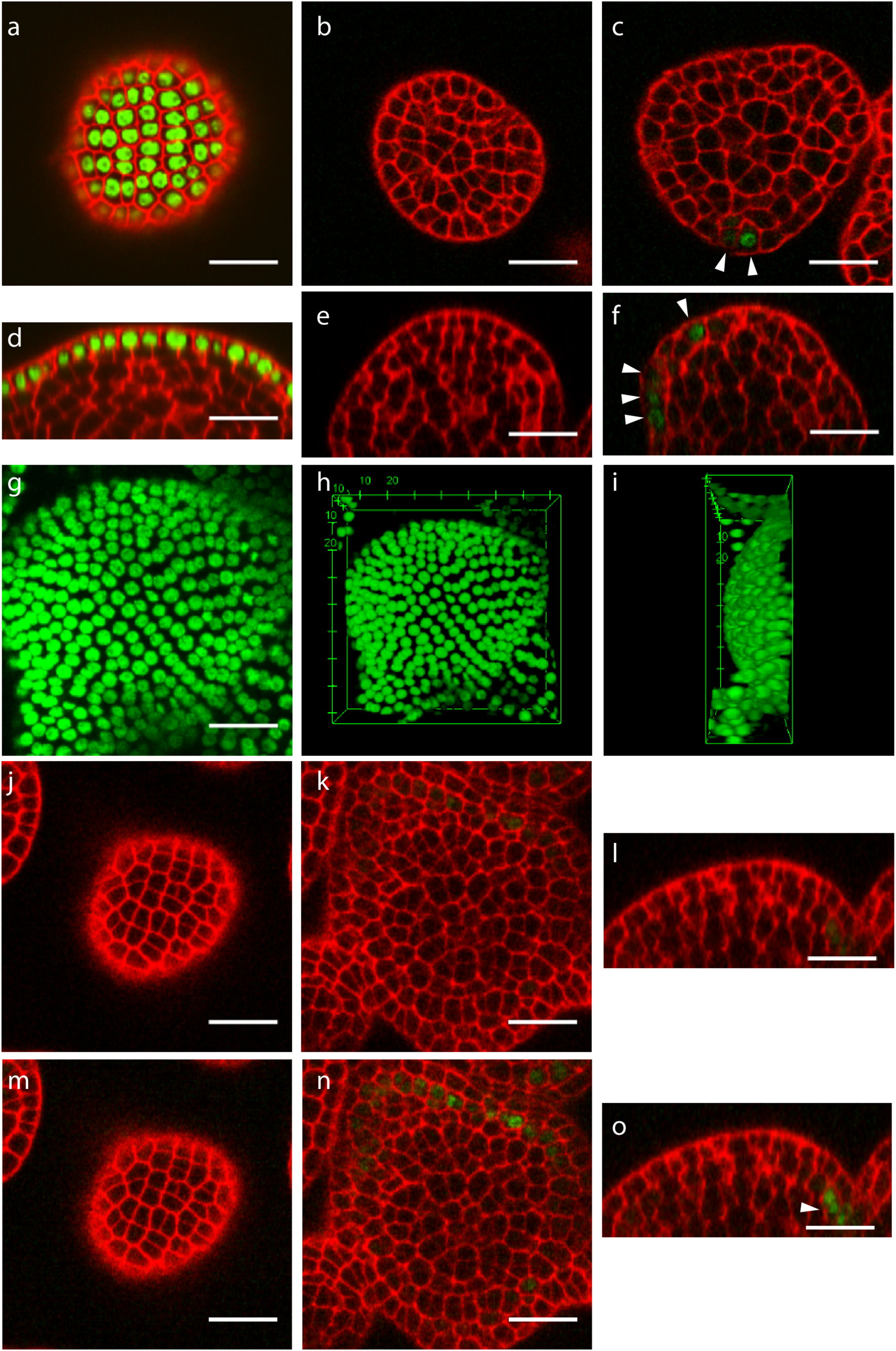
L1-specific gene editing of *pATML1:H2B-GFP* marker in the SAMs using a CRISPR-Cas9 system. **a, d**, Top (xy) (**a**) and orthogonal (xz) (**d**) view of two optical sections of a SAM from the parental line that expresses the *pATML1:H2B-GFP* marker. **b, e**, Top (xy) (**b**) and orthogonal (**e**) view (xz) of two optical sections of a flower primordium from a first-generation CRISPR mutation line, showing no GFP positive cells in L1. **c, f**, Top view (xy) (**c**) and orthogonal view (yz) (**f**) of two optical sections of another flower primordium from the same line as shown in (**b** and **e**), GFP positive cells are indicated (arrowhead). **g-i**, Maximum intensity projection of one SAM from a second-generation offspring of the founder line shown in (**b** and **c**). *pATML1:H2B-GFP* signal is normally expressed in whole SAM epidermis. **h, i**, Top (**h**) and side (**i**) 3D view of the same SAM shown in (**g**). 3D morphology of the SAM is normal. **j**, Top view of one xy optical section through the epidermis of a SAM from another second-generation offspring of the founder line shown in (**b-c**). None of cells shows GFP signal. **k**, **l**, Top (xy) (**k**) and orthogonal (yz) (**l**) view of optical sections through deep layers of the same SAM as shown in (**j**). **m-o**, Enhanced H2B-GFP signal intensity (multiplied by three times) as shown in (**j-l**) indicates a few cells, in L2, showing weak GFP-positive signal in (**n, o**). Arrowhead points to a group of cells with GFP positive signals in (**o**). **a-o**, Green color indicates H2B-GFP signal. Red color indicates cell walls that are stained with Propidium Iodide. scale bars = 20 µm.

We next used a GFP-fused translational reporter line. PIN1 protein is one of the proteins that mediates polar transport of auxin in the SAM, and the auxin maxima that result from PIN1 function specify the positions of new floral primordia^15^. In *pin1-/-* stable loss of function mutants, the inflorescence shoot apex becomes a naked pin-like structure with no functional flower primordium formation^16^. We used a *pPIN1:PIN1-mGFP5* reporter line in a *pin1-4* homozygous mutant background as the parental material for epidermally specific gene knockout^15^. Normal phyllotactic pattern is found in the SAMs of parental plants, and PIN1-GFP is expressed in future primordial sites in L1 and in the forming vascular tissues in underlying layers^17,18^ (Fig. 2a, d-g). We adopted the same CRISPR-Cas9 construct as used above to disrupt the PIN1-GFP protein sequence in the complementation reporter line (Fig. S1b). Since the GFP tag is inserted in the first exon of the *PIN1* gene (the predicted third cytoplasmic domain of the PIN1 transmembrane protein), a frame shift in the *GFP* coding sequence would affect both GFP fluorescence and the reading frame of the part of PIN1 encoded by the sequence downstream of *GFP*. In the T_1_ generation of the transgenic plants, we found SAMs with an epidermally depleted PIN-GFP fluorescence. Abnormal *pPIN1:PIN1-GFP* expression patterns were associated with SAMs that were surrounded by reduced numbers of irregularly arranged flowers (Fig. 2b-c). The mutated SAMs failed to initiate new floral primordia and showed naked pin-like shape that mimics the *pin* mutant phenotype (Fig. 2h-k). The older flowers seen in Fig. 2b-c were possibly formed at an early stage that, when SAMs were young and PIN1 protein was fully or partially functional, before gene mutagenesis happened in the entire targeted domain. The system thus apparently provides the possibility to generate materials to observe initial loss-of-function mutant phenotypes. The data are consistent with a previously reported result of selective removal of a PIN1-GFP transgene using a Cre-Lox system^19^. In the mutated SAMs, moderate intensity of GFP signals was still observed in L2 and in some corpus cells (Fig. 2j, k, Fig. S2). However, the presumptive provascular cells are arranged in a more random and diffuse pattern in these SAMs, suggesting that although PIN1 is still expressed in L2 of the mutated SAMs, it cannot instruct primordial initiation (as also indicated by the mutant phenotype) or vascular tissue formation (Fig. 2j, k). In total, we screened 105 T_1_ PIN1-GFP CRISPR transgenic plants, 12 of them (approximately 11%) show phyllotactic pattern defects and L1-specific suppression of PIN1-GFP fluorescent signal (Table S1). To track the mutation rate in the next generation, we screened progeny of one L1 mutant line in the T_2_ generation and found 4 of 32 (approximately 13%) of the plants showed a typical mutant phenotype and domain-specific fluorescent signal defect. In one of these four T_2_ plant SAMs, we found PIN1-GFP signal was diminished in multiple cell layers (the signal is only detected in corpus cells) (Fig. S3). This may be due to a CRISPR genome editing event at a very early embryo developmental stage before periderm formation, when the epidermis and sub-epidermis become separate clones. It is also possibly due to the low *ML1* promoter activity detected in L2 (Fig. S4) activating sufficient amounts of Cas9 transcription for mutational activity. PCR-based single clone sequencing also detected the somatic nature of CRISPR-induced target site specific mutations in the genomic DNA from three different lines that showing domain-specific PIN1-GFP signal suppression (Table S2). Line #3 corresponds to the line described above in which PIN1-GFP signal has multi-layer depletion (Table S2).

**Figure 2.**
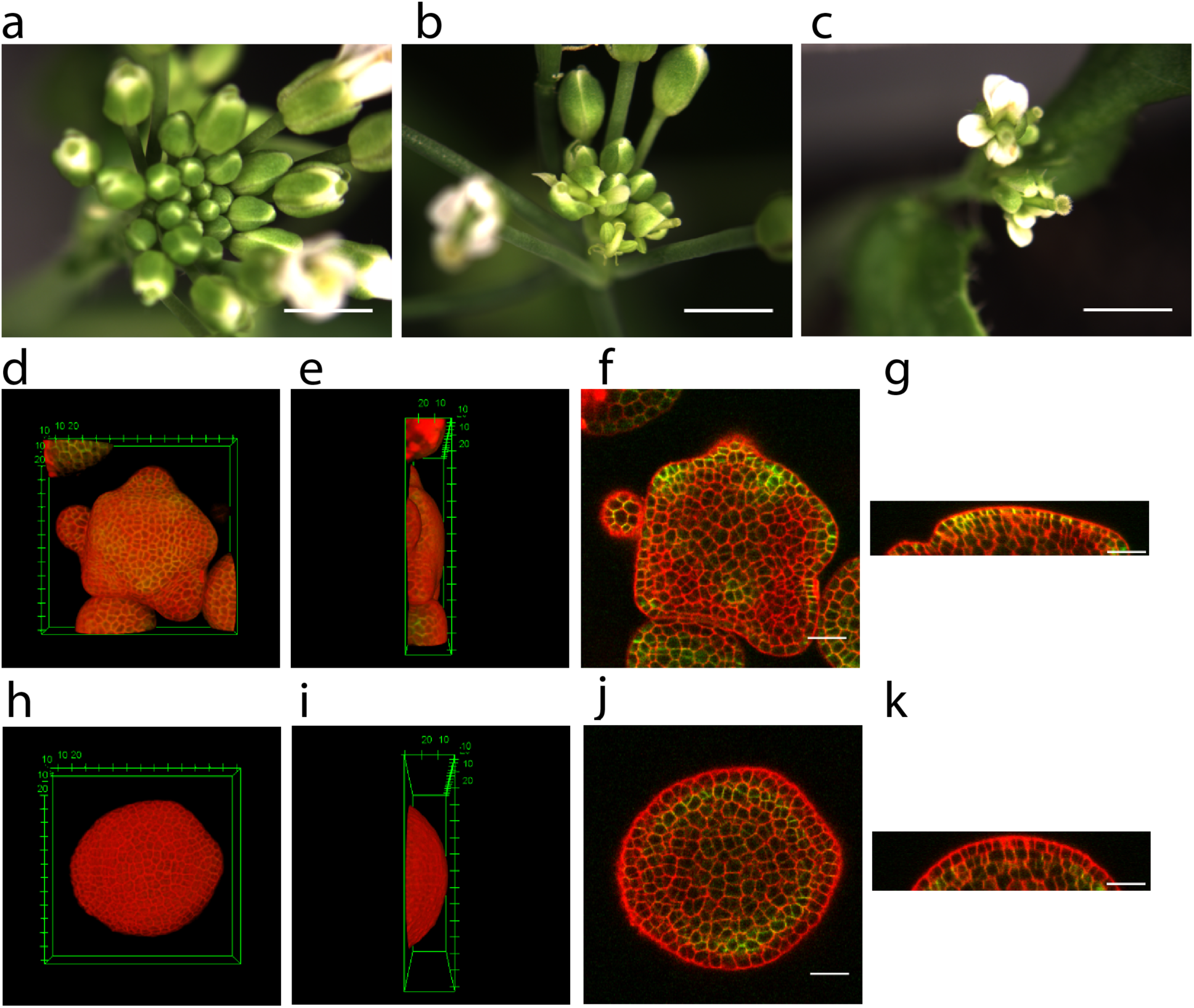
Epidermally specific gene editing of *pPIN1:PIN1-GFP* in the *Arabidopsis* shoot apex using the CRISPR-Cas9 system. **a**, A representative bright-field image of a parental shoot apex expressing the *pPIN1:PIN1-GFP* marker. **b, c**, Two representative bright-field images of shoot apices expressing the *pPIN1:PIN1-GFP* marker with *PIN1-GFP* knockout specifically in the epidermis. **d, e**, Top (D) and side (E) 3D view of a parental SAM expressing the *pPIN1:PIN1-GFP* marker (green). **f, g**, Top (xy) (**f**) and orthogonal (xz) (**g**) view of two optical sections through deep layers of the same SAM as shown in (**d**, **e**). 3D morphology of the SAM is normal and PIN1-GFP protein is enriched in the epidermis and presumed provascular tissue of the SAMs. **d-g**, Cell walls were stained with propidium iodide (red). **h, i**, Top (**h**) and side (**i**) 3D view of a SAM with *pPIN1:PIN1-GFP* marker in which the *PIN1-GFP* gene is specifically mutated in the epidermis. The SAM appears as a naked pin after older flowers are removed, as in a whole-plant *pin1* mutant. PIN1-GFP (green) is not visible in the epidermal layer in a 3D view, red indicates propidium iodide staining. **j, k**, Two optical sections through deep layers in top (xy) (**j**) and orthogonal (yz) (**k**) view of the same SAM as shown in (**h, i**). PIN1-GFP signal is maintained in subepidermal cells. **a-c**, Scale bars = 2 mm. **d-k**, 3D view image scale units are µm. 2D image scale bars = 20 µm.

Finally, we studied the possible mechanisms underlying the phyllotactic phenotype of L1-specific PIN1-GFP knockout lines. In the current model, PIN1 localization pattern is proposed to be determined by the mechanical stress level of cell walls, which directs auxin transport^20^. As a first hypothesis, we assumed that the mechanical sensor for PIN1 polarity control is absent in sub-epidermal and deeper layers, and therefore that the PIN1 protein expressed in L2 cannot perceive and respond to the mechanical stress field. As an alternative hypothesis, based on the model that considers the SAM as an inflated elastic thin shell^21^, we reasoned that in the L2 of the SAMs, mechanical stress differences are much smaller than in the epidermis because the majority of tensional forces are counterbalanced by the epidermal layer (which is under tension). In this scenario, L2-expressed PIN1 protein cannot form clear polarization patterns due to the lack of sufficient initial mechanical stress difference between different anticlinal cell walls, even though stress sensors are present, and PIN1 localization can respond to them. We carried out two different mechanical perturbation experiments to discriminate between the two hypotheses. First, osmotic treatment was applied to change plant cell mechanical properties^22,23^. We treated L1 mutated SAMs with 0.55 M mannitol solution to trigger plasmolysis and test the effect of turgor loss in PIN1-GFP localization in L2 cells. Previous studies showed that PIN1 protein undergoes rapid internalization upon turgor reduction^24^. After 90 minutes of mannitol treatment, PIN1 signal in L2 cells of L1-mutated SAMs was internalized (*n = 3* SAMs, Fig. 3a, b), suggesting the response of PIN1 protein to turgor perturbation is the same in the L2 of the SAMs as in L1. Secondly, we introduced strong mechanical perturbation by local cell ablations in both L1 and L2 cells of L1-mutated SAMs, which in the L1 causes a circumferential high stress pattern surrounding the ablation site^20^. In L2 cells of mutated SAMs, PIN1-GFP formed a partially outward pattern, which was consistent with the mechanical stress pattern induced by cell ablations in wild-type L1 (*n = 3*, Fig. 3c, d ^20^). These experiments indicate that the mechanical stress sensing components are present in both L1 and L2 cells of the SAMs, and that the cells in these layers both can respond to mechanical stress. The absence of an L1-like polarity pattern of PIN1 protein in the L2 of the mutated SAMs is therefore proposed to be due to reduced mechanical stress input, not to inability to sense mechanical stress.

**Figure 3.**
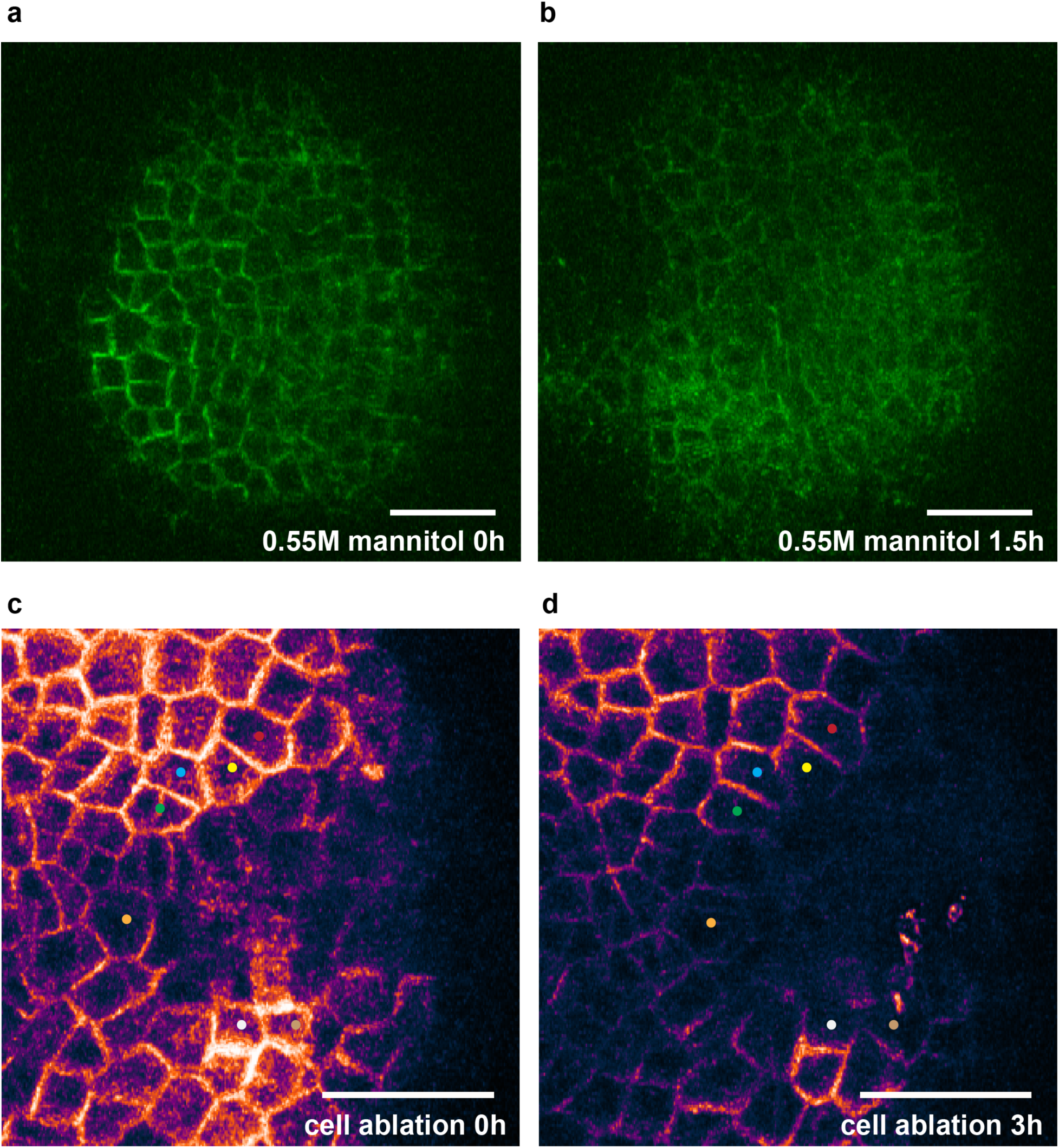
Test of mechanically induced PIN1-GFP protein relocalization in L2 cells. **a**, PIN1-GFP signal pattern, from L2 cells in a L1 *pin1-gfp* mutation line, just prior to 0.55M mannitol treatment. **b**, PIN1-GFP localization pattern, from the sample shown in (**a**), after 1.5 h mannitol treatment. PIN1-GFP protein was internalized after treatment. **c**, PIN1-GFP localization in L2 cells in a L1 *pin1-gfp* mutation line, immediately after local ablation of both L1 and L2 cells. **d**, PIN1-GFP protein partially polarized to the plasma membrane adjacent to walls away from the ablation site in surrounding cells at 3h after ablation, as compared with protein localization at 0h as shown in (**c**). Dots indicate representative cells in which PIN1-GFP protein relocalization has started to occur. Scale bar: 20 μm.

In summary, we present an efficient domain-specific genome editing approach through a combination of live imaging and a spatially-controlled CRISPR-Cas9 expression system in *Arabidopsis* shoot meristems. The high mutation rate (~20%) makes the system applicable to observe later effects of mutations that cause embryonic lethality or early developmental defects. Moreover, we demonstrated that the mechanical sensing that leads to subcellular PIN1 polarity exists in the L2 of the SAM, and that the abnormal phyllotaxis found in L1-specific PIN1-GFP knockout mutants is thus due to reduced mechanical stress differentials in the L2, and not to an inability of PIN1 in L2 cells to respond to stress. Live imaging-assisted expression domain specific CRISPR has potential application to improvement of domain-specific traits of crops, and to studies of *in vivo* gene function in post-embryonic developmental stages of animals and plants.

## Acknowledgments

We are grateful to Dr. Zachary L. Nimchuk for sharing the Nimchuk lab CRISPR system. We also thank Dr. Changfu Yao and Meyerowitz lab members for suggestions and discussions. The authors’ work was funded by the Howard Hughes Medical Institute and NASA grant NNX17AD53G to E.M.M..

## Contributions

T.L., A.Y., and E.M.M. conceived the experiments. T.L., A.Y. performed experiments. A.Y., T.L., and E.M.M. wrote the manuscript.

