## supplemental file for "Live imaging-assisted domain-specific CRISPR genome editing at single cell resolution in plants"

### METHODS

**Plant materials and growth conditions.** The *pATML1:H2B-mGFP*, *pPIN1:PIN1-mGFP5* were previously described [1,2]. *Arabidopsis thaliana* plants were grown in a sunshine soil/vermiculite/perlite mixture under continuous light at 20 °C.

**CRISPR-Cas9 system construction.** We created the tissue-specific CRISPR-Cas9 plant binary vector based on a *pHEE401E* backbone from the *Arabidopsis thaliana* egg-cell specific CRISPR-Cas9 system [3]. Specifically, we replaced its *Flag-Cas9-NLS* with *Arabidopsis* codon-optimized *Cas9-N7* coding sequences using *XbaI* and *SacI* restriction sites [4] and replaced its *U6-speR* with a *ccdB* gene flanked with *BsaI* restriction sites on both ends by use of the *HindIII* and *SpeI* restriction sites for Golden Gate cloning of multiple gRNA expression cassettes. We replaced the egg cell-specific promoter with the *ATML1* promoter (3.3kb, [1]) using *NcoI* and *SpeI* (the *SpeI* site was introduced upstream of the *Cas9* start codon by In-Fusion cloning), designated as *p1300-ccdB-ATML1p:AtCas9N7-hyg*. We adopted the *U6-26, U6-1, U3b* and *U3d* promoters [5,6] for gRNA expression. Between the RNA Polymerase III promoters and single guide RNA coding sequences without targeting sequences, using *HindIII* and *SpeI,* we inserted a *ccdB* gene flanked by *BsmBI* sites on both ends. The entire gRNA expression cassettes flanked with *BsaI* sites on both ends were ligated into the *pCR8 TOPO* vector, the result was named *pCR8-U6-26p-*

*ccdB-sgRNA*; *pCR8-U3bp-ccdB-sgRNA; pCR8-U6-1p-ccdB-sgRNA;* and *pCR8-U3dp-ccdB-sgRNA*. Oligonucleotides were synthetized based on targeting sequence, ligated into these four vectors by *BsmBI* digestion after annealing. The gRNA expression cassettes were ligated in a predefined order (*U6-26p:sgRNA, U3bp:sgRNA, U6-1p:sgRNA,* then *U3dp:sgRNA*) into the *p1300-ccdB-ATML1p:AtCas9N7-Hyg* vector by Golden Gate Cloning using *BsaI* digestion [7,8]. *p1300-gGFP-ATML1p:AtCas9N7-hyg* was then transformed into two *Arabidopsis thaliana* lines using the *Agrobacterium*-infiltration (floral dip) method [9] with hygromycin resistant $T_1$ plants used for further mutation screening. See Supplementary Table for the plasmid sequence information.

**Confocal imaging of fluorescent reporters in living tissues.** Plants bearing the *pATML1:H2B-mGFP*, *pPIN1:PIN1-mGFP5* fluorescent reporters were imaged using a Zeiss LSM 510 Meta confocal microscope or a Zeiss LSM 780 laser scanning confocal microscope. The laser and filter settings were as described previously [1,2]. To detect the L2 GFP signal in *pATML1:H2B-mGFP* fluorescent reporter, we increased the laser power from 3% to 6% when using a 488 nm excitation line and adjusted the gain setting from 750 to 800. Confocal z-stack images were processed using Fiji.

**Mechanical perturbation procedures.** For the osmotic treatment, we immersed dissected SAMs in 0.55M mannitol solution for 90 min. Imaging of PIN1-GFP signal was performed at 0h (right before treatment) and 1.5h (right after treatment). Due to light scattering by the mannitol solution, we imaged the samples in water. The cell ablations were performed manually with a pulled glass pipette under a Zeiss SteREO Discovery V8 stereomicroscope. Imaging of PIN1-

GFP signal was performed at 0h and 3h. The dissected SAMs were supported on growth medium agar plates (1 x MS salts + 1% sucrose + MES pH 5.7 + 0.8% Bacto Agar).

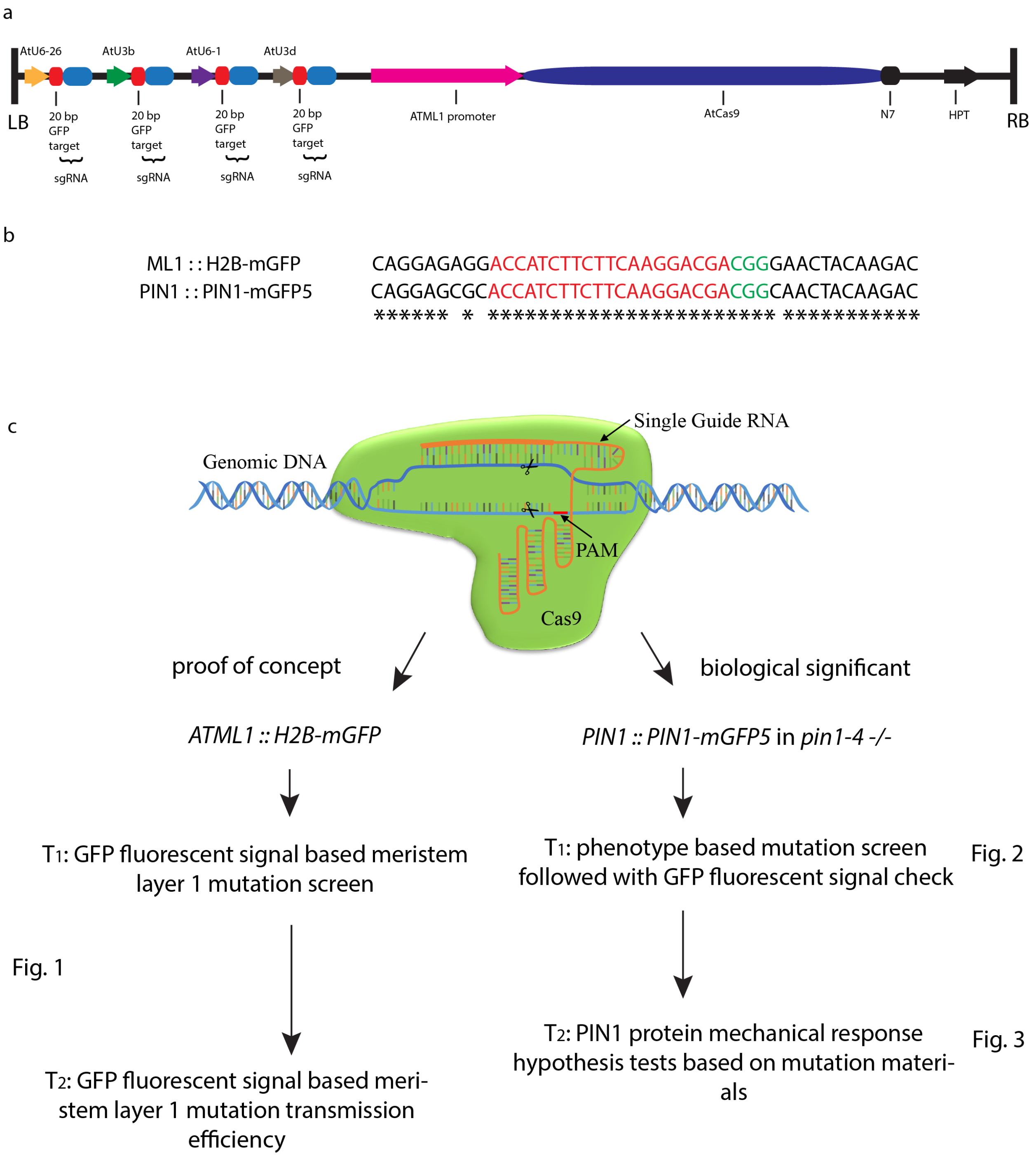
a
AtU6-26
AtU3b
AtU6-1
AtU3d
LB
20 bp GFP target
sgRNA
20 bp GFP target
sgRNA
20 bp GFP target
sgRNA
20 bp GFP target
sgRNA
ATML1 promoter
AtCas9
N7
HPT
RB
b
ML1 : : H2B-mGFP CAGGAGAGGACCATCTTCTTCAAGGACGACGGGAACTACAAGAC
PIN1 : : PIN1-mGFP5 CAGGAGCGCACCATCTTCTTCAAGGACGACGGCAACTACAAGAC
****** * *********************** ***********
c
Single Guide RNA
Genomic DNA
PAM
Cas9
proof of concept
biological significant
ATML1 :: H2B-mGFP
PIN1 :: PIN1-mGFP5 in pin1-4 -/-
T1: GFP fluorescent signal based meristem layer 1 mutation screen
T1: phenotype based mutation screen followed with GFP fluorescent signal check
Fig. 2
Fig. 1
T2: GFP fluorescent signal based meristem layer 1 mutation transmission efficiency
T2: PIN1 protein mechanical response hypothesis tests based on mutation materials
Fig. 3

**Supplementary Figure 1. Domain-specific CRISPR-Cas9 system information. a**, Schematic diagram of the sgRNAs and Cas9 expression cassettes in a plant binary vector (only elements between left border (LB) and right border (RB) that integrate into the plant genome are shown). **b**, Guide RNA targeting sequence (red colored DNA) information. Green colored DNA indicates PAM sequence. Asterisk marks the common sequence of *mGFP* and *mGFP5* after alignment. **c**, Schematic illustrating the work flow of this study.

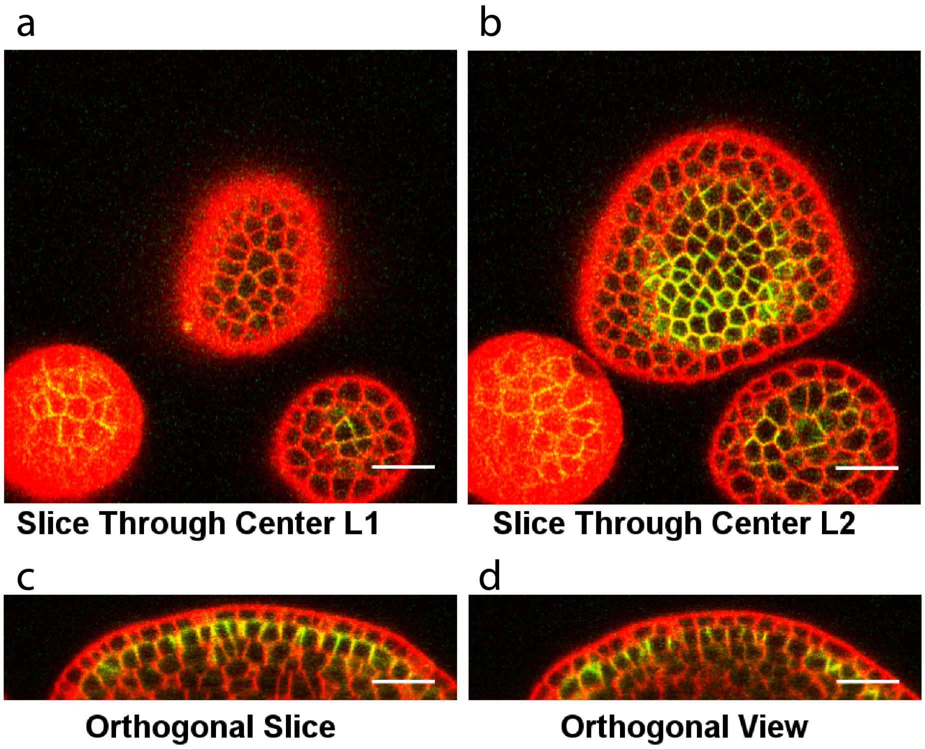


**Supplementary Figure 2**. **Visualization of PIN1-GFP signal in L2 in one mutated SAM. a**, A single horizontal optical section through the central region of L1. **b**, A single horizontal optical section through the central region of L2. **c-d**, two orthogonal optical section reconstructions of the same SAM. PIN1-GFP expression can be detected in every L2 cell (based on cell membrane signal). Scale bars = 20 μm.

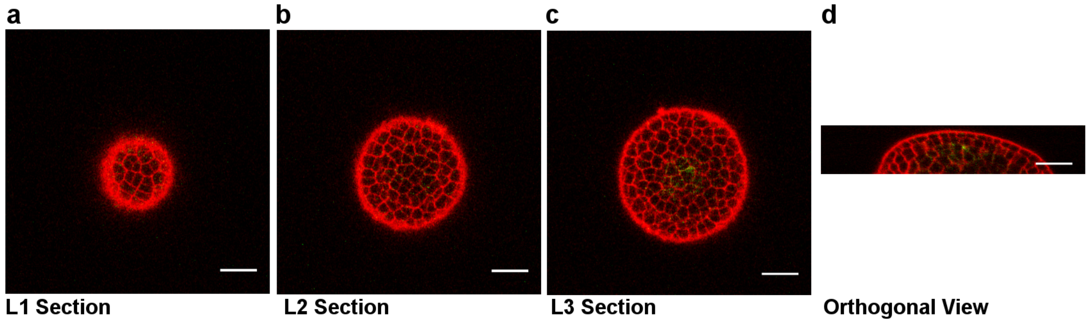


**Supplementary Figure 3**. **Visualization of PIN1-GFP signal in L3 in one mutated SAM.** **a**, A single horizontal optical section through the central region of L1. **b**, A single horizontal optical section through the central region of L2. **c,** A single horizontal optical section through the central region of the third cell layer. **d**, one reconstructed orthogonal section of the same SAM. PIN1-GFP expression can only be detected in corpus cells (based on cell membrane signal). PIN1-GFP was knocked out in first two layers of this particular SAM. Red: PI. Green: PIN1-GFP (the green signal was boosted 3-fold for better visualization). Scale bars = 20 μm.

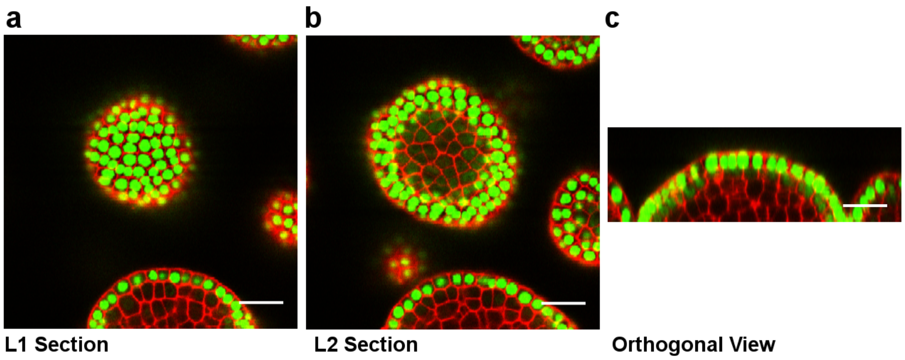


**Supplementary Figure 4**. ***ML1* promoter has low activity in L2 of the SAM. a**, A single horizontal optical section through the central region of the L1 of a SAM expressing *pML1::H2B-GFP*. **b**, A single horizontal optical section through the central region of the L2 of the same SAM as in (A). **c**, one orthogonal reconstruction of a section through the same SAM as in (**a**). Red: PI. Green: H2B-GFP. The GFP signal in this image stack was purposely extremely over-exposed to reveal the low-level activity of the ML1 promoter in the L2 layer. Faint H2B-GFP signals are present in the L2 layer as shown in (**b-c**). Scale bars = 20 μm.

**Supplementary Table 1: Gene targeting frequencies at *GFP***

| Plant generations | *pATML1::H2B-mGFP*[1,2] | *pPIN1::PIN1-mGFP5*[1,2] |
|---|---|---|
| $T_1$ | 19/90 (21%) | 12/105 (11%) |
| $T_2$ | 6/15 (40%) | 4/32 (13%) |

1. Values are shown as number of SAMs that contain a fluorescent signal defect in epidermis / total number of SAMs that were screened (gene targeting frequency %). The frequency is rounded to the nearest percent.
2. In both $T_1$ and $T_2$ generations, fluorescent signal-based screening was used for identification of *pATML1::H2B-mGFP* tissue-specific mutations. Phenotypic screening combined with fluorescent signal confirmation was used for PIN1::PIN1-mGFP5 tissue-specific mutation identification.

**Supplementary Table 2. Molecular basis for mutated targeting site**

| mutation line from $T_2$ generation | targeting sequence | mutation type | mutated ratio[1] |
|---|---|---|---|
| Wildtype | ACCATCTTCTTCAAGGACGACGG | NA | 0/48 (0%) |
| *pATML1::H2B-GFP* #1 | ACCATCTTCTTCA---------- | -42bp | 2/40 (5%) |
| *pATML1::H2B-GFP* #2 | ACCATCTTCTTCAAGGA-GACGG<br>ACCATCTTCTTCAAGGAgCGACGG | -1bp<br>+1bp | 2/39 (5%)<br>1/39 (3%) |
| *pATML1::H2B-GFP* #3 | ACCATCTTCTTCAAGGA-GACGG | -1bp | 2/40 (5%) |
| *pATML1::H2B-GFP* #4-1[2] | ACCATCT---------ACGACGG<br>ACCATCTTCTTCAAGtccgccatgcc | -9bp<br>+22bp | 1/48 (2%)<br>1/48 (2%) |
| *pATML1::H2B-GFP* #4-2[2] | ACCATCTT-------gtaGACGG | -10/+3bp | 1/48 (5%) |
| *pATML1::H2B-GFP* #5 | ACCATCTTCTTCAAGGACGACGG | NA | 0/48 (0%) |
| *pATML1::H2B-GFP* #6 | -------------------<br>ACCATCTTCTTCAAGGAaCGACGG<br>ACCATCTTCTT------------ | -25bp<br>+1bp<br>-12bp | 2/31 (6%)<br>1/31 (3%)<br>1/31 (3%) |
| *pPIN1::PIN1-GFP* #1 | ACCATCTTCTTCAAGGAtCGACGG | +1bp | 8/34 (24%) |
| *pPIN1::PIN1-GFP* #2 | ACCATCTTCTTCAAGGAgCGACGG | +1bp | 2/23 (9%) |
| *pPIN1::PIN1-GFP* #3 | ACCATCTTCTTCAAGGAaCGACGG<br>ACCATCTTCTTCAAGGAtCGACGG | +1bp<br>+1bp | 12/25 (48%)<br>5/25 (20%) |

1. Values are shown as number of clones containing mutated allele / total number of examined clones (gene targeting frequency %). The frequency is rounded to the nearest percent.
2. *pATML1::H2B-GFP* #4-1 and #4-2 samples were from the same mutation line but from two different axillary meristems.

**Supplementary Table 3.** Plasmid sequences for experiments in this study.

| Plasmid Name | Link to plasmid sequences in Benchling |
|---|---|
| p1300-gGFP-ATML1p:AtCas9N7-hyg | https://benchling.com/s/seq-fjry7hYD5NwUIEWDofl4 |
| p1300-ccdB-ATML1p:AtCas9N7-hyg | https://benchling.com/s/seq-FWOYBxXS4DYys9GMG32V |
| pCR8-U6-26p-ccdB-sgRNA | https://benchling.com/s/seq-McF0JTxOWdvdu59evyEH |
| pCR8-U3bp-ccdB-sgRNA | https://benchling.com/s/seq-7UFcbDQMTmyelzA7s5sO |
| pCR8-U6-1p-ccdB-sgRNA | https://benchling.com/s/seq-VoDULYFS36pDLSbp1duU |
| pCR8-U3dp-ccdB-sgRNA | https://benchling.com/s/seq-AmuYjiSIpxMNfTTRhSYS |
